# Analysis of Large Peptide Collisional Cross Section Dataset Reveals Structural Origin of Bimodal Behavior

**DOI:** 10.1101/2025.09.10.675470

**Authors:** Allyn M. Xu, Dániel Szöllősi, Helmut Grubmüller, Oded Regev

## Abstract

Recent advances in ion mobility spectrometry have enabled the measurement of rotationally averaged collisional cross-sectional area (CCS) for millions of peptides, as part of routine proteomic mass spectrometry workflows. One of the most striking finding in recent large ion mobility datasets is that CCS exhibits two distinct modes, most notably for charge 3^+^ peptides. Here, using classical machine learning approaches, we identify that basic site positioning is a key sequence feature determining if a peptide belongs to the high or low CCS mode. Molecular dynamics simulations suggest that peptides in the high CCS mode tend to adopt more extended conformations and form charge-stabilized helical structures, whereas those in the low CCS mode adopt more compact, globular conformations. Further supporting this structural hypothesis, we provide evidence for preferential protonation near the C-terminus, and uncover multiple position-dependent sequence determinants that all suggest the predominance of helix formation in the high mode. Together, these findings will enable better integration of CCS measurements into protein identification and quantification pipelines, improving the performance of ion mobility-based proteomics.

## Introduction

Ion mobility spectrometry (IMS) is a powerful, widely used technique for measuring the size and shape of gas-phase analytes with applications to many fields, including structural biology,^1^ proteomics,^2–4^ and more.^5^ In IMS, ions under the influence of an electric field are separated based on collisions with buffer gas,^6,7^ enabling measurement of the rotationally averaged collisional cross-sectional area (CCS), thereby providing insights into the conformation of analytes. IMS has been used in structural studies of biomolecules, complementing other methods such as crystallography, cryo-electron microscopy, or NMR spectroscopy.^1,8,9^ More recently, IMS paired with high-throughput mass spectrometry has enabled enhanced separation and analysis of complex analyte mixtures.^2–4^

IMS has also been extensively used to study the conformations of polypeptides in the gas phase.^1,10,11^ Studying peptides in the gas phase can be informative for many reasons. The gas phase provides a simplified environment for studying intramolecular forces due to the absence of solvent interactions.^10,11^ Moreover, the gas phase may be a more representative environment than the solution phase for certain biological environments, such as in the interior of the cell membrane, which has a low dielectric constant of *ϵ* ≈ 3.^12^ There are some notable differences in the formation of secondary structure for peptides in the gas phase. Specifically, various studies, employing IMS combined with molecular dynamics (MD) simulations, have shown that *α*-helices more readily form when in alignment with the peptide’s charges.^13–18^ For example, the peptide ion Ac–Ala_*n*_–LysH^+^ (*n* = 10–20) displays a CCS consistent with an *α*-helical structure, whereas Ac–LysH^+^–Ala_*n*_ displays a lower CCS consistent with a compact, globular structure.^13,15^ This difference is attributed to the fact that the C-terminal proton interacts favorably with the negative end of the *α*-helix’s macrodipole, which arises from the arrangement of the peptide backbone’s N–H and C=O functional groups.^19^ On the other hand, the N-terminal charge destabilizes the *α*-helix, and instead the charge is solvated directly by the carbonyl groups. Similar studies have found evidence for other compact conformations, such as a folded-helix, depending on the position of charges.^14,16^ Despite these insights, these studies are restricted to peptides composed mostly of alanines, an amino acid with notably high helix propensity,^20^ and there still remain many unanswered questions about gas-phase conformations of peptides in general.

Recently published large-scale datasets have provided a general outlook on CCS across millions of peptides.^21,22^ In these works, a single CCS is assigned to each peptide, corresponding to the most intense measured peak (even though some peptides can exhibit multiple stable conformations^23^). Interestingly, when CCS is plotted against mass, two distinct CCS modes appear, most prominently seen for charge 3^+^ peptides (Fig. 1). It was noted that this bimodality shares similarities with the extended and compact conformations observed in the previous polyalanine *α*-helix studies.^21,22^ However, it is still not fully understood how a peptide’s sequence influences its propensity to adopt the low or high CCS mode, or more generally, its CCS. In principle, this could depend on a complex interplay of structure, intramolecular forces, and the dynamics of gas-phase charges.^24–28^

**Figure 1.**
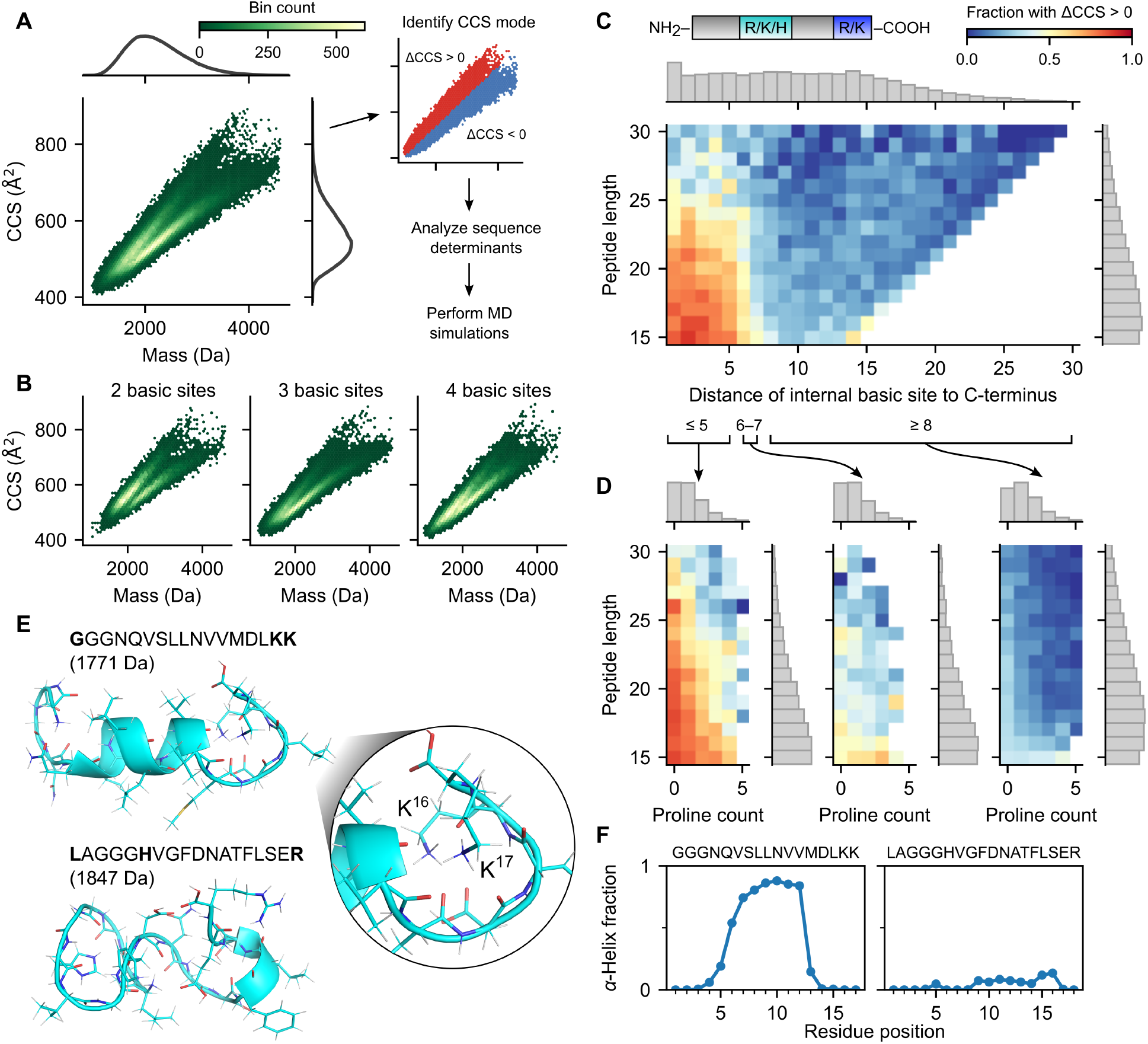
CCS bimodality stratified by select peptide features. **(A)** 2D hexagonal binning plot of CCS versus mass for charge 3^+^ peptides in Meier et al.’s dataset, with marginal densities for CCS and mass shown on right and top axes, respectively (left panel). Analysis workflow, including same binning plot in the left panel, but colored red and blue for the low and high CCS mode, respectively (right panel). **(B)** 2D hexagonal binning plot of CCS versus mass for charge 3^+^ peptides with two basic sites (left), three basic sites (middle), and four basic sites (right). **(C)** 2D histogram of peptide length versus distance (in amino acids) of internal basic site to C-terminus for charge 3^+^ non-Nt-acetylated tryptic peptides with three basic sites, colored by fraction of peptides in bin with ΔCCS *>* 0 (difference between CCS and CCS_separator_; see methods). Marginal density histograms on right and top axes, respectively (grey boxes). Cartoon of peptide shown at the top, with internal basic site colored in teal, and C-terminal arginine or lysine colored in blue. **(D)** 2D histogram of peptide length versus proline count, with coloring and marginal histograms as in panel (C), for peptides with an internal basic site with distance to C-terminus less than or equal to 5 (left), between 6 and 7 (middle), and greater than or equal to 8 (right). Bins with less than 8 peptides were excluded. **(E)** Backbone ribbon and side chains stick representation of MD simulation for representative three basic site peptides from the high and low CCS modes, respectively. Basic sites are shown in bold. **(F)** Plot of the fraction of MD simulation runs where a given residue was involved in an *α*-helix, for each of the two representative peptides.

Here, we analyze a large-scale CCS dataset^22^ and perform MD simulations to investigate structural explanations for the observed bimodality. We find that peptides having at least two basic sites in the C-terminal region strongly favor the high CCS mode whereas those with one or no basic sites favor the low CCS mode. MD simulations demonstrate that in the former case, two C-terminal protonated basic sites form a cap stabilizing a *α*-helix, providing a structural explanation for the high CCS mode. MD simulations also show that in the latter case, peptides tend to form compact, globular conformations, explaining the low mode. To further support this helix formation hypothesis, we use interpretable machine learning models that reveal the position-dependent effects of amino acids. These models provide evidence that residues near the C-terminal end are preferentially protonated, consistent with helix macrodipole effects. They also uncover (1) a preference in high CCS mode peptides for hydrophobic amino acids in the region expected to form a helix, as well as (2) a helix-breaking effect of glycine in high (but not low) CCS mode peptides, an effect we confirm with additional MD simulations. All these findings are physicochemically consistent with the helix formation hypothesis. The insights provided here into peptide gas-phase conformations and their sequence determinants should inform future ion mobility-based proteomics pipelines.

## Methods

### Data collection

A post-processed CCS dataset was obtained from Meier et al.^22^ As detailed in their work, peptides were generated from a variety of organisms (*C. elegans, D. melanogaster, E. coli, H. sapiens*, and *S. cerevisiae*) using three proteases (trypsin, LysC, and LysN), processed through fractionated LC-TIMS-MS/MS runs on a TIMS-quadrupole TOF mass spectrometer, and analyzed using the MaxQuant software. For each peptide and charge state, a single CCS was selected per LC-TIMS-MS run based on the most abundant identified feature. Finally, each peptide and charge state was assigned a unique CCS value, obtained from aligning and averaging across LC-TIMS-MS runs.

The dataset contains CCS values for 559,979 unique combinations of peptides and charge states. For this study, we focused on the 143,850 peptides with charge 3^+^. The peptide lengths range from 8 to 55 amino acids, with a median of 19.

### Separating peptides based on low and high CCS modes

To identify the low and high modes of the CCS bimodality (Fig. 1a), we first fit a Gaussian mixture model (*n* = 2) on the 2D dataset of CCS versus mass. Classifying the points based on the Gaussian mixture model, we observed that the separator was nearly linear. For simplicity, we then fit a linear separator on the Gaussian mixture model classification using squared hinge loss (regularization parameter = 1.0). The resulting linear separator is given by

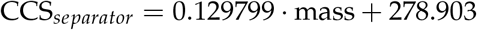

where CCS is given in Å^2^ and mass in Da.

For each peptide, we computed the difference between its CCS and CCS_*separator*_, referred to as ΔCCS. Lastly, a peptide was denoted as belonging to the low or high CCS mode if ΔCCS ≤ 0 or ΔCCS *>* 0, respectively.

### Predicting CCS mode and absolute CCS using generalized additive models

We trained generalized additive models (GAMs)^29^ to investigate the sequence determinants of both CCS mode and absolute CCS. Specifically, we trained four G AMs: two logistic GAMs to predict CCS mode, and two linear GAMs to predict absolute CCS. The input to all models was the peptide sequence (as a string of amino acids), and the dependent variable was either CCS mode (coded as 0 for the low mode and 1 for the high mode) or absolute CCS (in Å^2^).

The four GAMs were trained on four different subsets of peptides from Meier et al.’s training dataset,^22^ filtered to include unmodified tryptic peptides with charge 3^+^, exactly three basic sites, and a minimum length of 15 residues. For CCS mode prediction, one model was trained on peptides where the internal basic site was located within five residues of the C-terminus, and another on peptides where the internal basic site was at least seven residues away. For absolute CCS prediction, each of these two subsets was further restricted to peptides in the high or low CCS mode, respectively, and a separate model was trained on each.

GAM predictions are based on the total contribution of: (1) the identity and relative position of each non-C-terminal amino acid in the peptide sequence, (2) the peptide length in residues, and (3) the identity of the C-terminal amino acid (R or K).

We designed a specialized “shared spline” architecture to enable the GAMs to operate on sequential inputs and capture position-specific contributions of amino acids. Each of the 20 standard amino acids is assigned a learned spline function (denoted *f*_*a*_ for amino acid *a*). For a peptide sequence *a*_1_*a*_2_ … *a*_*n*_, the contribution of the *i*th non-C-terminal amino acid *a*_*i*_ is given by its corresponding spline function 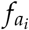, evaluated at the amino acid’s relative position within the peptide. This relative position is given by 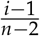, ranging linearly from 0 (N-terminal) to 1 (penultimate residue). In other words, the spline *f*_*a*_ encodes the contribution that amino acid *a* as a function of its relative position.

The contribution of peptide length is captured by a learned spline *f*_len_, and the contribution of the C-terminal amino acid is given by an indicator variable 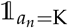—equal to 0 if the residue is arginine (R), and 1 if lysine (K)—multiplied by a learned coefficient *β*_C-term. K_.

Together, the total contribution of a peptide input sequence *a*_1_*a*_2_ … *a*_*n*_ is:

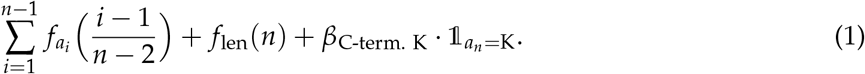

For CCS mode GAMs, the output is the standard logistic function of the total contribution above, interpreted as the predicted probability that the peptide belongs to the high CCS mode. For the absolute CCS GAMs, the total contribution is interpreted directly as the predicted CCS value (in Å^2^).

The models were implemented in Python using the pyGAM library,^30^ with custom modifications to support the shared spline architecture. Hyperparameters—including the number of splines and smoothing penalties on derivative and magnitude—were tuned to balance model fit and spline smoothness. Code and trained models are linked in the “Code availability” section.

### Molecular Dynamics Simulations

In order to sample the conformational space of the peptide sequences in question, we performed fully atomistic, *in vacuo* molecular dynamics temperature quenching simulations. Two sets of sequences were tested (P1: **GGG**NQVSLLNVVMDLKK, P2: **GGG**LAHVGFDNATFLSER), each with the position of the glycine triplet progressively shifted toward the C-terminus, resulting in 14 and 15 peptides tested, respectively. Fully extended conformations were generated using Pymol (Version 3.0 Schrödinger, LLC.) to ensure unbiased starting structures.

All peptides carried a charge of 3^+^, assigned to the basic amino acids and the N-terminus. Acidic amino acids were protonated and thus neutral. After topology generation, peptides were placed in a cubic box with a side length of 100 nm. Energy minimization was performed using the steepest descent integrator with 3000 steps, followed by a four-step equilibration procedure with increasing time steps of 0.1 fs, 0.5 fs and 1 fs with position restraints. The first three setups were conducted under an NVT ensemble and the final under an NPT ensemble. The resulting structures served as the starting point for the quenching simulations.

A simulated annealing temperature quench was applied to allow the peptides to reach an equilibrium conformation. Specifically, the initial temperature was set to 600 K and maintained for 10 ns, then linearly decreased to 305 K over a time period of 0.5 *μ*s and subsequently kept at 305 K for another 40 ns to facilitate thermal equilibration.

The net charge of the system and the absence of solvent required a specific set of simulation parameters: (i) GROMACS (version 2023.4)^31^ was compiled in double precision and run with a shorter than usual 1 fs integration time step; (ii) cutoff distances were set to 30 nm, such that all peptide atoms interacted with all other atoms explicitly via Coulomb and Lennard-Jones forces at all times, allowing a long neighbor search interval (1 ns) and therefore, enhancing performance; (iii) Particle Mesh Ewald (PME) method was not used, as it would fail given the total net charge of the system; (iv) pressure coupling was disabled. The system was periodic, and the center of mass was kept at the center of the simulation box. The temperature was controlled by the V-rescale algorithm.^32^

All simulations were performed with the CHARMM36m force field.^33^ The quenching procedure was repeated 1000 times for each peptide, with starting velocities randomly drawn from a Boltzmann distribution, separately for each run. All atomic positions were saved every 10 ns.

## Results & Discussion

### CCS bimodality of peptides with three basic sites is largely determined by internal basic site position

Charge 3^+^ peptides in Meier et al.’s dataset display a bimodal distribution when CCS is plotted versus mass (Fig. 1A). We therefore separated the peptides based on whether they exhibit CCS in the low or high mode (see Methods). We found that approximately 60% of these peptides belong to the low mode, while 40% belong to the high mode, and this bimodality persists even when considering peptides with a specific number of basic sites (arginine, lysine, histidine, or N-terminus amine group) (Fig. 1B). These two modes are largely distinct and well-separated, except in the lower mass ranges. To ensure that only peptides with well-defined modes were considered, we limited our analysis to those with 15 or more amino acids. Furthermore, we restricted our analysis to tryptic and LysN peptides that were not N-terminal acetylated (Nt-acetylated), which made up the majority (98%) of the dataset (see methods for details). For these 120,466 peptides (mass range: 1370–4600 Da), we investigated the sequence features that explain their CCS bimodality.

To minimize ambiguity in charge localization, we first focused on the 60,396 peptides with exactly three basic sites. For such peptides, the 3^+^ charge is likely due to protonation of all three basic sites. Tryptic peptides in this subset have fixed basic sites at the N- and C-termini, with one additional internal basic site positioned anywhere along the peptide. In contrast, LysN peptides in this subset have two basic sites fixed at the N-terminus, while a third basic site can occur anywhere.

Interestingly, for tryptic peptides, we observed a distinct transition between the two CCS modes depending on the position of the internal basic site (Fig. 1C). Specifically, among tryptic peptides with an internal basic site located near the C-terminus (within five amino acids), 68% exhibited CCS in the high mode, whereas among tryptic peptides with an internal basic site farther from the C-terminus (at least seven amino acids), 79% exhibited CCS in the low mode. This trend was most pronounced in peptides between 15 and 25 amino acids in length, while it was diminished in peptides longer than 25 amino acids. With their basic sites located further towards the N-terminus, LysN peptides extended this trend, showing a similar percentage of 81% in the low mode when the (sole) internal basic site is near the C-terminus, and 99% otherwise (Supplementary Fig. S1). Together, these findings demonstrate that basic sites in the proximity of the C-terminus promote the high CCS mode—markedly so when two such sites are present.

One explanation for these observations is that protonated basic sites located near the C-terminus may induce charge-stabilized helical-like structures, leading to high CCS values. Specifically, such sites can interact favorably with backbone carbonyl groups that, in a helical conformation, are exposed toward the C-terminal end. Similar structures have been observed in molecular dynamics simulations, suggesting their viability as candidate conformers.^16^ In contrast, protonated basic sites located farther from the C-terminus can destabilize helical structures due to unfavorable electrostatic interactions with the helix macrodipole. This can result in more compact, globular conformations that exhibit low CCS values.^14^

This reasoning aligns with that presented by Counterman et al.,^14^ whose MD simulations showed that charge 3^+^ polyalanines favor extended helical conformations when charges are predominantly located in the C-terminal half, and compact conformations when charges are predominantly located in the N-terminal half. Our observations suggest that similar structural principles apply much more broadly to a wide range of peptides. We note that Counterman et al. focused on the relative position of charges along the peptide (i.e., as a fraction of peptide length), whereas our findings suggest that it is the absolute position (specifically, having two basic amino acids within 5–7 residues of the C-terminus) that plays a more critical role. Indeed, we observed little evidence that relative position matters beyond absolute position (Fig. 1C, Supplementary Fig. S1).

To further support the hypothesis that the high CCS mode largely consists of charge-stabilized helical-like structures, we evaluated the effect of proline and length—two features we expected to disrupt such conformations. Namely, proline is known to break helical structures,^34^ while longer peptides exhibit greater conformational freedom, which we reasoned may make it more difficult to maintain stable helical conformations. Indeed, among tryptic peptides with three basic sites, we observed that the fraction of peptides in the high CCS mode decreased with both increasing proline count and increasing peptide length, even when stratified by the internal basic site position (Fig. 1D). These results are consistent with the interpretation that the high CCS mode corresponds to extended, helix-like conformers.

To provide additional support for the helical hypothesis, we performed MD simulations on two charge 3^+^ peptides with three basic sites: one from the high CCS mode with the internal basic site adjacent to the C-terminal lysine (GGGNQVSLLNVVMDLKK; CCS = 515 Å^2^, ΔCCS = 6.7 Å^2^) and the other from the low CCS mode with the internal basic site closer to the N-terminus (LAGGGHVGFDNATFLSER; CCS = 490.7 Å^2^, ΔCCS = –27.9 Å^2^). In the simulations, the high mode peptide frequently formed a helical conformation (Fig. 1E, top; Fig. 1F, left; Supplementary Fig. S5A), with the helix capped at the C-terminal end by two lysine charges. These two lysine charges were solvated by the backbone carbonyl groups of the helix and of the capping structure (Fig. 1E, right). This structural arrangement supports the hypothesis that extended, helix-like conformations can be stabilized by electrostatic interactions between C-terminally localized charges and the helix macrodipole. In contrast, the low mode peptide adopted compact, globular conformations with minimal secondary structure (Fig. 1E, bottom; Fig. 1F, right; Supplementary Fig. S5B), consistent with the expectation that basic sites located away from the C-terminus destabilize helices.

In summary, our results show that peptides with three basic sites tend to adopt the high CCS mode when two of those sites are located near the C-terminus. This supports a model in which gas-phase peptide conformation reflects a balance between helix-promoting and helix-disrupting features, with C-terminal charge localization favoring extended, helical-like structures. The additional effects of proline and peptide length, together with simulation-based evidence of charge-stabilized helices, further reinforce this framework. Our findings align with prior studies on polyalanine and suggest that charge-stabilized helical conformations may be a general structural motif across a wide range of peptide sequences.

### Position of second but not first internal basic site affects CCS mode of peptides with four basic sites

To further elucidate the CCS bimodality, we next examined charge 3^+^ peptides with more than three basic sites. Specifically, we examined non-Nt-acetylated tryptic peptides with four basic sites, again restricting our analysis to peptides that are at least 15 amino acids long. These peptides contain an amine group at the N-terminus, an arginine or lysine at the C-terminus, and two internal basic sites whose positions vary across the peptides. Note that with four basic sites, the localization of the 3^+^ net charge is clearly ambiguous.

Interestingly, we found that CCS mode strongly correlated with the position of the second internal basic site, and is largely independent of the position of the first. Specifically, peptides tended to adopt the high CCS mode when the second internal basic site was located within 8–10 residues of the C-terminus, and the low CCS mode otherwise (Fig. 2A). One explanation for this preference is that an internal basic site near the C-terminus enables a configuration in which two of the three protons can localize toward the C-terminal end—a distribution we hypothesized promotes the high CCS mode by facilitating charge-stabilized helix formation. This would explain why the high CCS mode is largely observed only when the second internal basic site is positioned sufficiently close to the C-terminus. In contrast, when no internal basic site is positioned near the C-terminus, such a charge distribution might not be possible. This may explain why these peptides predominantly fall into the low CCS mode.

**Figure 2.**
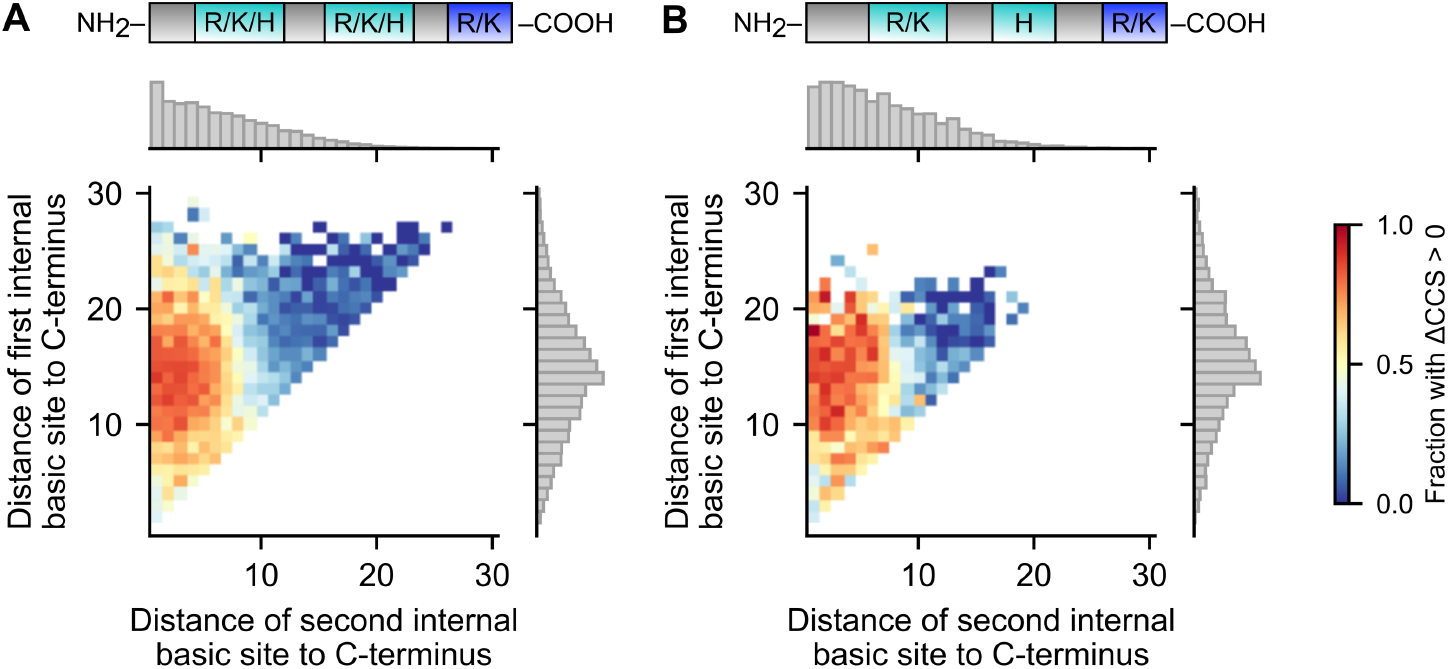
CCS bimodality in peptides with four basic sites. 2D histogram of the distance of the first versus second internal basic site from the C-terminus for charge 3^+^ non-Nt-acetylated tryptic peptides with four basic sites, colored by fraction of peptides in bin with ΔCCS *>* 0. Bins with less than 8 peptides were excluded. Marginal density histograms on right and top axes, respectively (grey boxes). **(A)** All analyzed peptides. **(B)** Peptides whose internal basic sites are an arginine or lysine followed by a histidine. Cartoon of peptide shown at the top, with internal basic sites colored in teal, and C-terminal arginine or lysine colored in blue.

Interestingly, we observed that this positional effect persists even when the first internal basic site is an arginine or a lysine and the second is a histidine (Fig. 2B). The side chains of arginine and lysine are more basic than that of histidine, whose gas-phase basicity is generally comparable to that of the N-terminal amine.^35,36^ Given that the four basic sites compete for three protons, one would not expect histidine to be a favored site of protonation in such peptides. However, our findings suggest that histidine is frequently protonated when positioned near the C-terminal end. This is consistent with the idea that charge localization is influenced not only by the basicity of the side chains but also by their spatial alignment with structural features—particularly the macrodipole associated with a helical-like conformation. Indeed, prior studies have shown that macrodipole effects can shift the apparent basicity or acidity of functional groups along the peptide backbone, favoring charge retention near the C-terminus.^37^

Furthermore, a similar pattern emerged even in peptides with more than four basic sites. Specifically, in charge 3^+^ tryptic peptides with five basic sites (N-terminal amine, three internal basic residues, and C-terminal R/K), the CCS mode largely depended on whether the last internal basic site was positioned near the C-terminus—regardless of the positions of the other two internal basic sites (Supplementary Fig. S2). Consistent with the reasoning above, this supports the notion that a helix may form when the protonation pattern enables stabilization by C-terminal charges, and conversely, that a C-terminal basic site is preferentially protonated when it aligns with a helical conformation, even when multiple sites compete for charges.

Based on these findings, we speculate that helix-like conformations and the associated macrodipole interactions may play a key role in guiding charge localization in the gas phase. Moreover, our results point to a feedback relationship: charge distribution can influence peptide structure, and structural features—such as alignment with the macrodipole—can, in turn, influence where charges are stabilized. This interplay offers a plausible mechanistic explanation for the observed CCS bi-modality, in which peptides with basic sites clustered near the C-terminus tend to favor extended, helix-like conformations.

### Sequence features influencing CCS mode

Despite the overall importance of internal basic site positioning in determining CCS mode, other sequence features also have an effect. Indeed, even when the internal basic site is near the C-terminus, some peptides are in the low mode, and conversely, even when the internal basic site is far from the C-terminus, some peptides are in the high mode (Fig. 1C). To systematically investigate these effects, we trained generalized additive models (GAMs)^29^ to predict CCS mode (see Methods). Each GAM prediction is modeled as the sum of position-dependent contributions from individual residues. Specifically, the model includes one learned spline for each of the 20 amino acids, capturing how residue contributions vary with their relative positions within peptides. This design allows us to capture potential amino acid effects that depend on their sequence position. Additionally, we included a spline function to model peptide length effects, and a categorical variable to account for the identity of the C-terminal amino acid (R or K).

We focused our analysis on unmodified tryptic peptides with three basic sites and a minimum length of 15 residues, similar to the set analyzed in the first results section. Given that the internal basic site’s position strongly influenced CCS mode (Fig. 1C), we stratified peptides into two sub-groups: those where the internal site was within five residues of the C-terminus, and those where it was more than seven residues away. We then trained a separate GAM for each subgroup, achieving high prediction accuracy (AUROC = 0.883 and 0.840, respectively).

First, we analyzed the learned spline functions for peptides in the subgroup where the internal basic site was near the C-terminus (Fig. 3A, red). As described above, we hypothesize that peptides in this subgroup tend to adopt extended *α*-helical structures stabilized by a pair of charges near the C-terminus (Fig. 1). Interestingly, the amino acid splines exhibit position-dependent contributions that are generally more pronounced in a broad central region, slightly skewed toward the C-terminus (Fig. 3A). These patterns suggest that this region plays a structurally relevant role in determining CCS mode, potentially indicative of structural elements such as *α*-helices.

**Figure 3.**
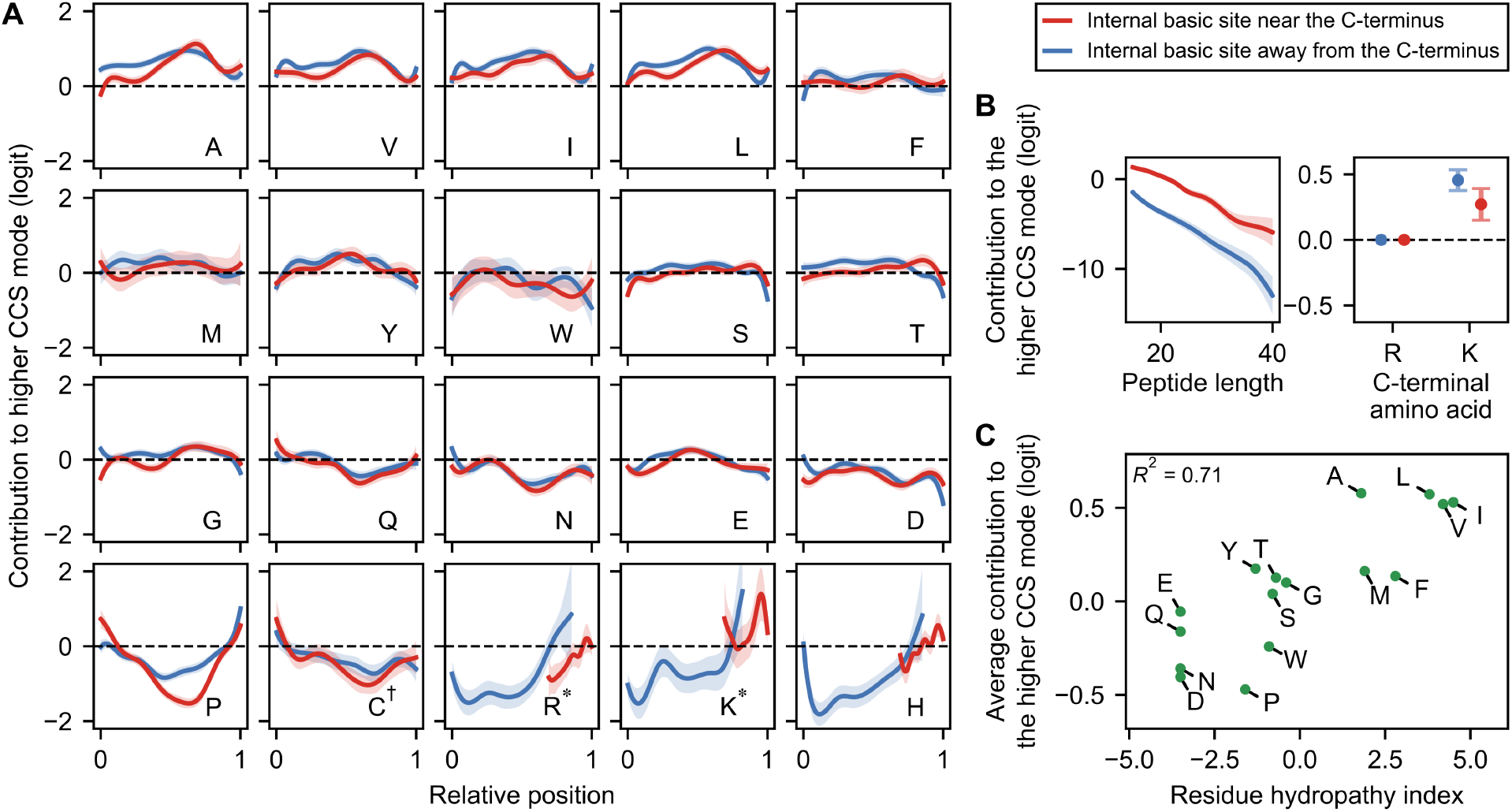
Logistic GAM models predicting CCS mode. **(A**,**B)** GAM models trained to predict CCS mode. Shaded regions indicate 95% confidence intervals. Each amino acid (except the C-terminal one) contributes additively to the final prediction on the logit scale, depending on its relative position along the peptide, as shown in the corresponding panel (A). The additive logit contribution of length and of the C-terminal amino acid are shown in (B). C^†^ refers to carbamidomethylated cysteine; R^*^, K^*^ refer to non-C-terminal arginine and lysine, respectively. **(C)** The contribution of each amino acid, averaged over all positions and over the two modes, as a function of its Kyte-Doolittle hydropathy. Carbamidomethylated cysteine is excluded.

Notably, amino acids with non-aromatic hydrocarbon side chains (A, V, I, L) exhibit similar spline patterns, generally favoring the high CCS mode when located in this central region (Fig. 3A). In line with our hypothesis, hydrocarbon side chains may be less disruptive to helix formation in the gas phase, due to their non-polar nature, which may explain their observed propensity to favor the high CCS mode. Indeed, valine has been shown to exhibit high *α*-helix propensity in the gas phase, in contrast to its behavior in aqueous solution.^38^ In fact, more broadly, spline contributions generally correlate with amino acid hydrophobicity (Fig. 3C): hydrophobic residues promote the high CCS mode, whereas hydrophilic residues such as Q, N, E, and D tend to decrease this likelihood. While hydrophobicity is not a direct proxy for gas-phase polarity, the observed correlation is consistent with the notion that residues with polar side chains—often classified as hydrophilic—may hinder *α*-helix formation in the gas phase through electrostatic disruption.^39^ In addition, proline exhibited the most pronounced spline pattern, substantially reducing the likelihood of adopting the high CCS mode when centrally located in the peptide (Fig. 3A). This aligns closely with its established role as a helix breaker,^34^ and is in agreement with our earlier observations (Fig. 1C). Lastly, peptides terminating in lysine favored the high CCS mode more strongly than those ending in arginine (Fig.3C). This may result from lysine’s smaller, more flexible side chain—potentially better suited to helix-capping—than the bulkier, planar guanidinium group of arginine. Collectively, these observations support the physicochemical relevance of the GAM outputs and align well with the hypothesis of charge-stabilized *α*-helical conformations.

Second, we examined the learned spline functions for peptides whose internal basic site was far from the C-terminus (Fig. 3A, blue). Notably, their spline patterns closely resemble those of the previous subgroup—where the internal basic site was near the C-terminus—suggesting that peptides in this subgroup may also adopt similar charge-stabilized helical conformations. We speculate that these conformations might involve two positive charges near the C-terminus (even though there is only one basic site there), with either the internal basic site or the N-terminal amine being unprotonated. Although speculative, this scenario is plausible: our results above suggest that charge localization near the C-terminus may be favorable due to potential alignment with an *α*-helix macrodipole (Fig. 2). Consistent with this idea, one of the strongest contributions for promoting the high CCS mode in this peptide subgroup occurs when the internal basic site is at the N-terminus and is a histidine (but much less so when it is an arginine or a lysine) (Fig. 3A; Supplementary Fig. S3). One plausible explanation for this pattern is that strong Coulombic repulsion at the N-terminus, combined with histidine having lower basicity compared to arginine and lysine, could make double protonation of the N-terminal amine and histidine less favorable energetically. This would promote localization of one of those charges near the C-terminal end, again potentially stabilizing an extended helix-like conformation. In summary, the formation of charge-stabilized helices provides a coherent explanation for the high CCS mode in this subgroup.

Together, our GAM analysis reveals sequence features consistent with the hypothesis that extended, charge-stabilized helical conformations underlie the high CCS mode. The position-specific influences of amino acid residues—especially the propensity of hydrophobic residues to promote helices, the destabilizing effects of helix-breaking residues like proline, and the differential effectiveness of lysine and arginine in C-terminal helix capping—all lend strong plausibility to this structural model.

### Sequence features influencing absolute CCS within each mode

Having examined how sequence features influence CCS mode, we next investigated how they determine CCS within each mode. As before, we used the set of unmodified tryptic peptides with exactly three basic sites and length at least 15 residues. We then divided these peptides into high and low CCS mode subgroups, training a linear GAM on each to predict absolute CCS (see Methods). These models allowed us to examine how specific residues and sequence patterns contribute to shifts in absolute CCS, rather than transitions between distinct classes of conformational states.

The GAMs capture intuitive relationships: CCS increases with peptide length in both subgroups (Fig. 4B), consistent with the physical intuition that longer peptides generally occupy more space. Additionally, the average value of the amino acid splines correlates strongly with residue mass (*R*^2^ = 0.83; Fig. 4C), reiterating the mass-CCS correlation observed earlier (Fig. 1A).

**Figure 4.**
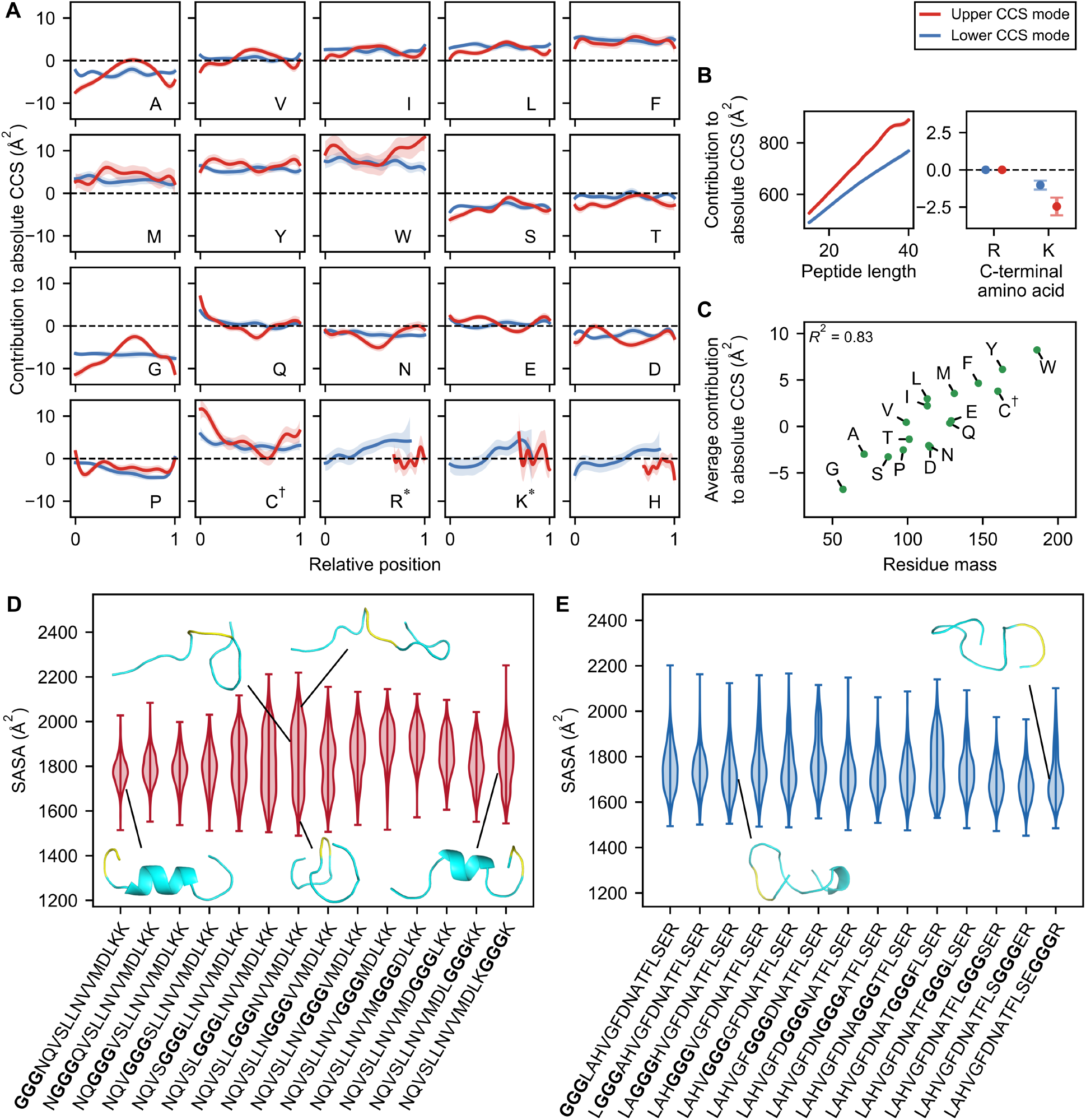
Linear GAM models predicting absolute CCS. **(A**,**B)** GAM models trained to predict absolute CCS. Shaded regions indicate 95% confidence intervals. Each amino acid (except the C-terminal one) contributes additively to the final prediction depending on its relative position along the peptide, as shown in the corresponding panel (A). The additive contribution of length and of the C-terminal amino acid are shown in (B). C^†^ refers to carbamidomethylated cysteine; R^*^, K^*^ refer to non-C-terminal arginine and lysine, respectively. **(C)** The contribution of each amino acid, averaged over all positions and over the two modes, as a function of its residue mass. **(D**,**E)** Molecular dynamics simulations of the high mode series (D) and the low mode series (E). Also shown are representative conformations. Glycine triplet is highlighted in yellow.

Although the learned splines from both GAMs exhibit similar average values (Pearson’s *r* = 0.98 across amino acids), their position-dependent patterns differ notably. In particular, the amino acid splines for the low mode subgroup are generally flat (Fig. 4A, blue); that is, each amino acid residue contributes similarly to CCS regardless of their position. This aligns with the notion that the low mode subgroup primarily comprises random, globular conformations, where residues influence CCS predominantly through their size rather than their specific positions. In contrast, amino acid splines for the high mode subgroup shows positional dependence (Fig. 4A, red). For instance, tryptophan increases CCS by ~7 Å^2^ when located closer to the peptide termini compared to a central position. A possible explanation is that in *α*-helical conformations, tryptophan’s bulky side chain is more exposed near the peptide termini, whereas the helix may shield it when it is positioned more centrally.

Surprisingly, glycine also exhibits pronounced position dependence: in the high mode subgroup, glycine increased CCS by ~10 Å^2^ when centrally located, compared to positions near the peptide termini (Fig. 4B, red). To further investigate this positional effect of glycine, we turned to MD simulations. We selected a peptide from the high mode subgroup containing a glycine triplet (GGGNQVSLLNVVMDLKK), then generated a series of peptides differing only by the position of this glycine triplet, by sliding it across consecutive positions. We then ran simulations for each peptide in this series, computing the solvent accessible surface area (SASA) as a proxy for CCS (see Methods). We observed that the peptides with the glycine triplet located in the center of the peptide indeed tended to exhibit larger conformations than those with the triplet located at the ends (Fig. 4D). Examining representative conformations, we found that when the glycine triplet was located near the termini, peptides formed *α*-helical conformations capped by the double lysine at the C-terminus (Fig. 4D, lower left and right diagrams; Fig. 1E). However, when the glycine triplet was located in the center, the peptides tended to form elongated, extended conformations, with low *α*-helical content (Fig. 4D, upper diagrams; Supplementary Fig. S4A, S5A). These simulations indicate that glycine may differentially influence extended conformers, by introducing local flexibility that perturbs helical structure when positioned at the center of the peptide. Interestingly, the presence of a glycine triplet towards the center of the peptide also induced larger variance in the simulated SASA values, due to the presence of globular conformations with lower SASA values (Fig. 4D, center diagram). This aligns with the notion that glycine can also destabilize extended conformations and promote compact conformers.

Lastly, we verified the observed glycine spline for the low CCS mode, by repeating the MD simulations, but for a different series generated using a peptide with a glycine triplet from the low mode subgroup (LAGGGHVGFDNATFLSER). Indeed, we observed that the simulated SASA values did not vary strongly with the position of the glycine triplet (Fig. 4E), and that the *α*-helical content was consistently low (Supplementary Fig. S4B, S5B). Moreover, we observed that representative conformations for these peptides were compact globular peptides, consistent with our hypothesis (Fig. 4E, diagrams).

Taken together, these results highlight how amino acid identity and sequence position jointly influence peptide CCS within conformational modes. Our GAM and MD analyses underscore how subtle positional effects can significantly modulate peptide structure.

## Summary & Conclusion

In this study, we investigated sequence determinants underlying the CCS bimodality in charge 3^+^ peptides. We found that CCS mode is strongly influenced by basic site positioning: peptides with basic sites distributed toward the C-terminus strongly favor the high mode. Together with our MD simulations, these findings support the hypothesis that such peptides often adopt extended helix-containing conformations. This extends previous observations from polyalanine systems and suggests that *α*-helical structures are prevalent in the gas phase across a broad range of peptides. We also found evidence consistent with the hypothesis that residues near the C-terminus are preferentially protonated due to helix macrodipole effects. These insights may inform MD simulations of gas-phase proton dynamics. Lastly, our GAM analyses revealed additional sequence features associated with CCS mode and absolute CCS. These sequence features correlate with the physicochemical properties of peptides and offer additional insights into the conformational landscape underlying peptide CCS.

Our study has a few limitations. First, the peptide ions in the large-scale dataset of Meier et al.^22^ were each assigned the strongest CCS peak, even though some peptides can exhibit other secondary CCS peaks.^23^ However, peak CCS has been demonstrated to be extremely consistent (an accuracy of *<*1%),^22^ suggesting that it provides a robust measure for a peptide’s preferred gas-phase conformation. Second, our analysis is influenced by the distribution of peptides in the selected dataset. Our use of a dataset spanning the proteomes of multiple organisms partly mitigates this issue. This can be further addressed by including peptides generated with more enzymes or by including synthetic peptides. Third, because to our knowledge no biomolecular force field for in vacuo simulations is available,^40^ we used a force field that has been parameterized using experimental data and MD simulations in solution. As a result, our simulations may be less accurate than those in solution; based on previous experience,^40^ however, we think they are sufficiently accurate to support the qualitative conclusions presented here.

In conclusion, our analysis has demonstrated sequence determinants that can largely explain the CCS bimodality for a wide-range of peptides. These findings provide insights that inform our understanding of the gas-phase conformations of peptides and the dynamics of their charges.

## Acknowledgements

We thank Jürgen Cox for his assistance with technical queries.

## Code availability

Code, trained models, and figure generation scripts are available on GitHub: https://github.com/regev-lab/peptide-ccs-analysis.

## Author contributions

A.X. and O.R.: Conceptualization and writing original draft. A.X.: Data analysis. D.S.: MD simulations. H.G. and O.R.: Supervision. A.X., D.S., H.G., and O.R.: Manuscript review, editing, and revision.

## Supplementary Figures

**Figure S1.**
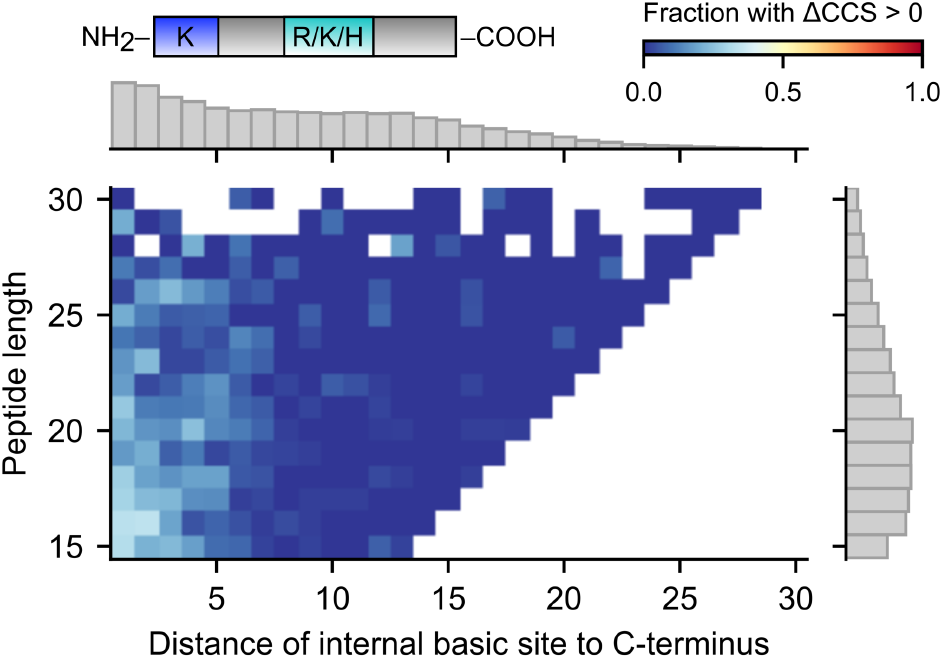
CCS bimodality for LysN peptides (related to Fig. 1C). 2D histogram of peptide length versus distance (in amino acids) of internal basic site to C-terminus for charge 3^+^ non-Nt-acetylated LysN peptides with three basic sites, colored by fraction of peptides in bin with ΔCCS *>* 0. Bins with less than 8 peptides were excluded. Marginal density histograms on right and top axes, respectively (grey boxes). Cartoon of peptide shown at the top, with internal basic site colored in teal, and N-terminal lysine colored in blue.

**Figure S2.**
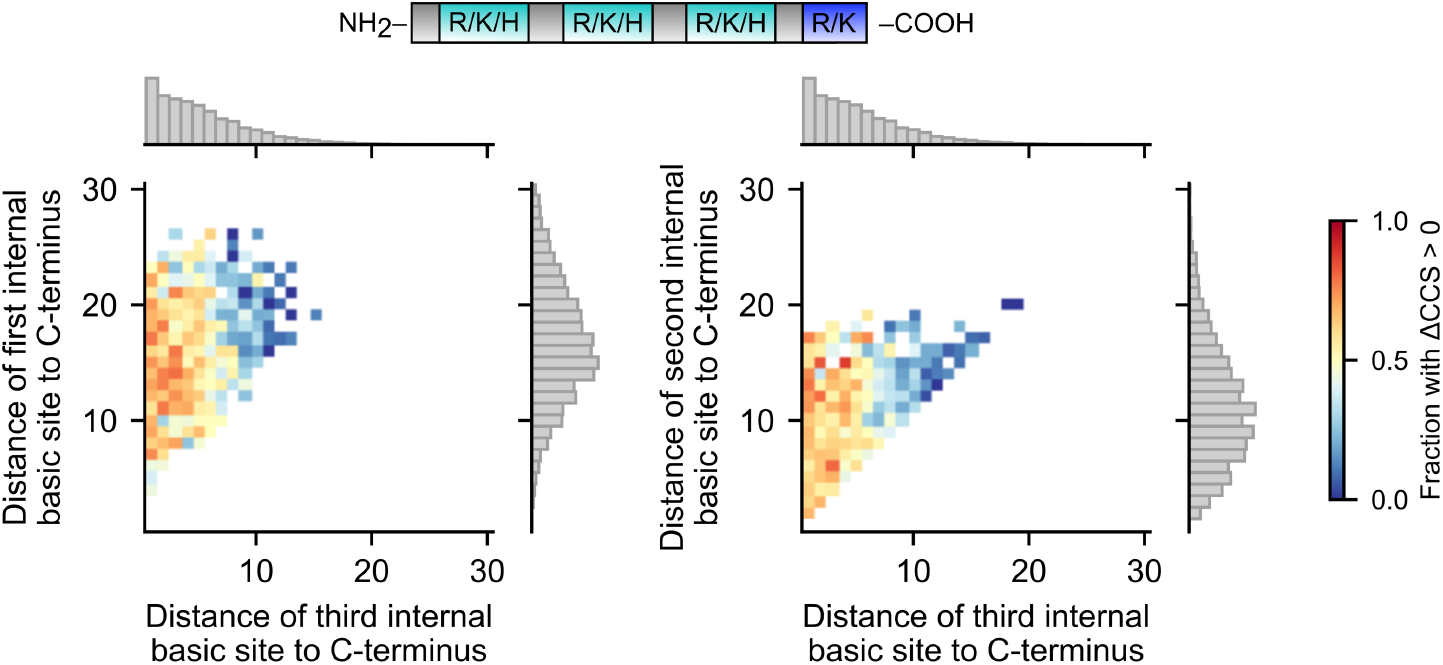
CCS bimodality in peptides with five basic sites (related to Fig. 2). 2D histogram of the distance of the first versus third internal basic site (left) and of the second versus third internal basic site (right) from the C-terminus for charge 3^+^ non-Nt-acetylated tryptic peptides with five basic sites, colored by fraction of peptides in bin with ΔCCS *>* 0. Bins with less than 8 peptides were excluded. Marginal density histograms on right and top axes, respectively (grey boxes). Cartoon of peptide shown at the top, with internal basic site colored in teal, and C-terminal arginine or lysine colored in blue.

**Figure S3.**
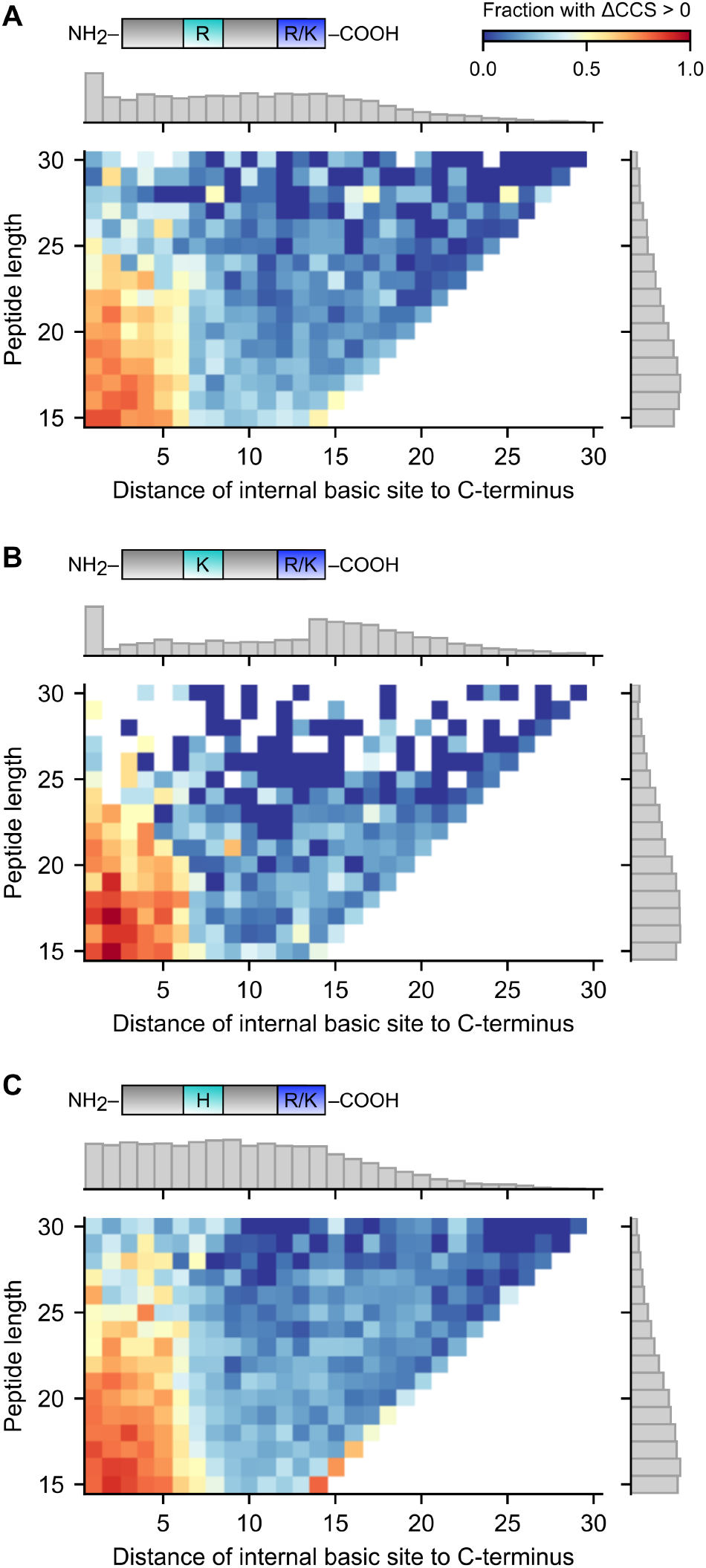
CCS bimodality with a specific internal basic site (related to Fig. 1C). 2D histogram of peptide length versus distance (in amino acids) of internal basic site to C-terminus for charge 3^+^ non-Nt-acetylated tryptic peptides with three basic sites whose internal basic site is a arginine **(A)**, lysine **(B)**, and histidine **(C)**, colored by fraction of peptides in bin with ΔCCS *>* 0. Right-most diagonal corresponds to the internal basic site being at the N-terminus. Bins with less than 3 peptides were excluded. Marginal density histograms on right and top axes, respectively (grey boxes). Cartoon of peptide shown at the top, with internal basic site colored in teal, and C-terminal arginine or lysine colored in blue.

**Figure S4.**
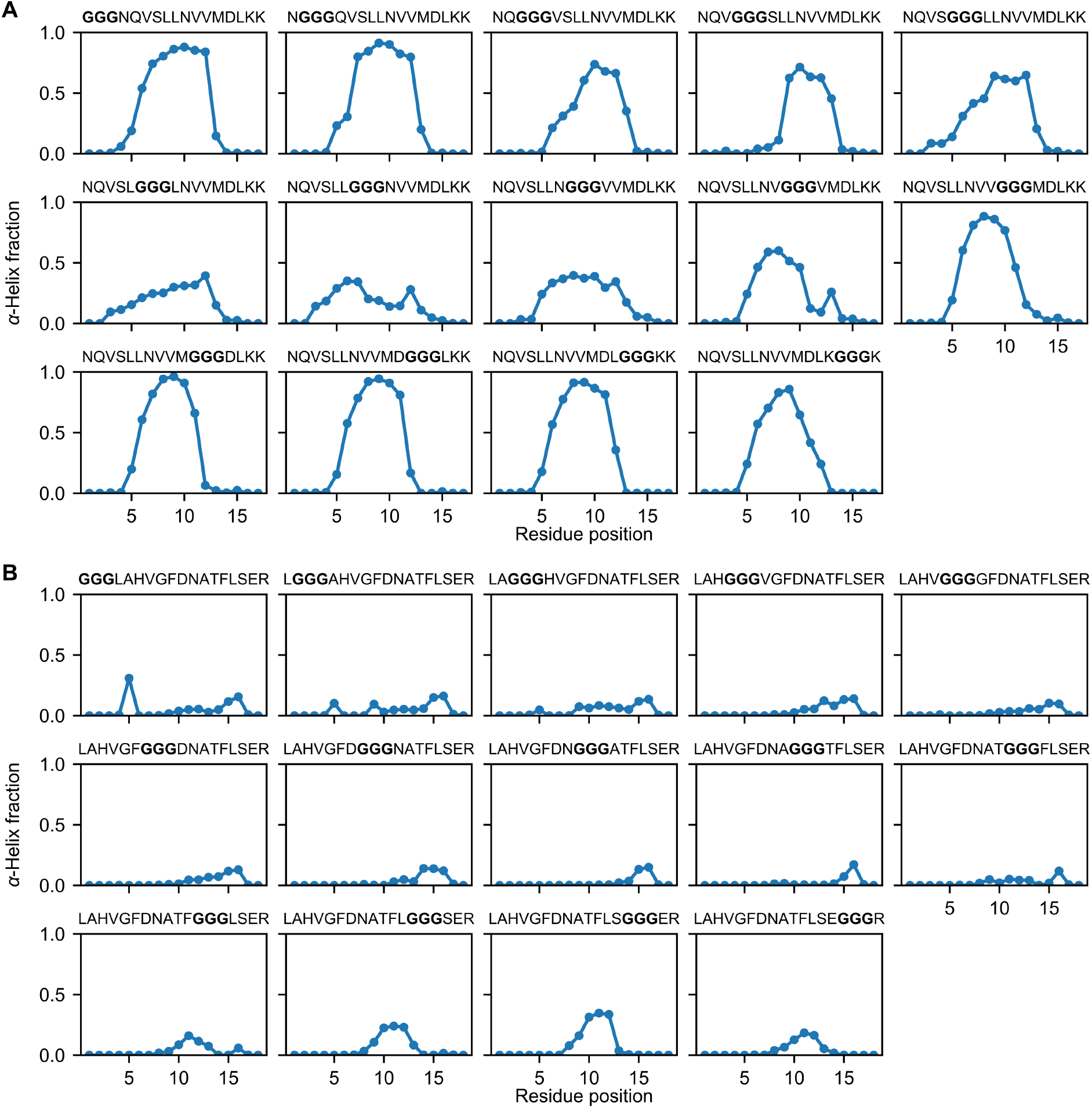
*α*-helix fraction by residue. Plot showing the fraction of MD simulation runs where a given residue was involved in an *α*-helix, for each simulated peptide. Plots corresponding to peptides in the high mode series grouped in (A) and the low mode in (B). Glycine triplet is shown in bold.

**Figure S5.**
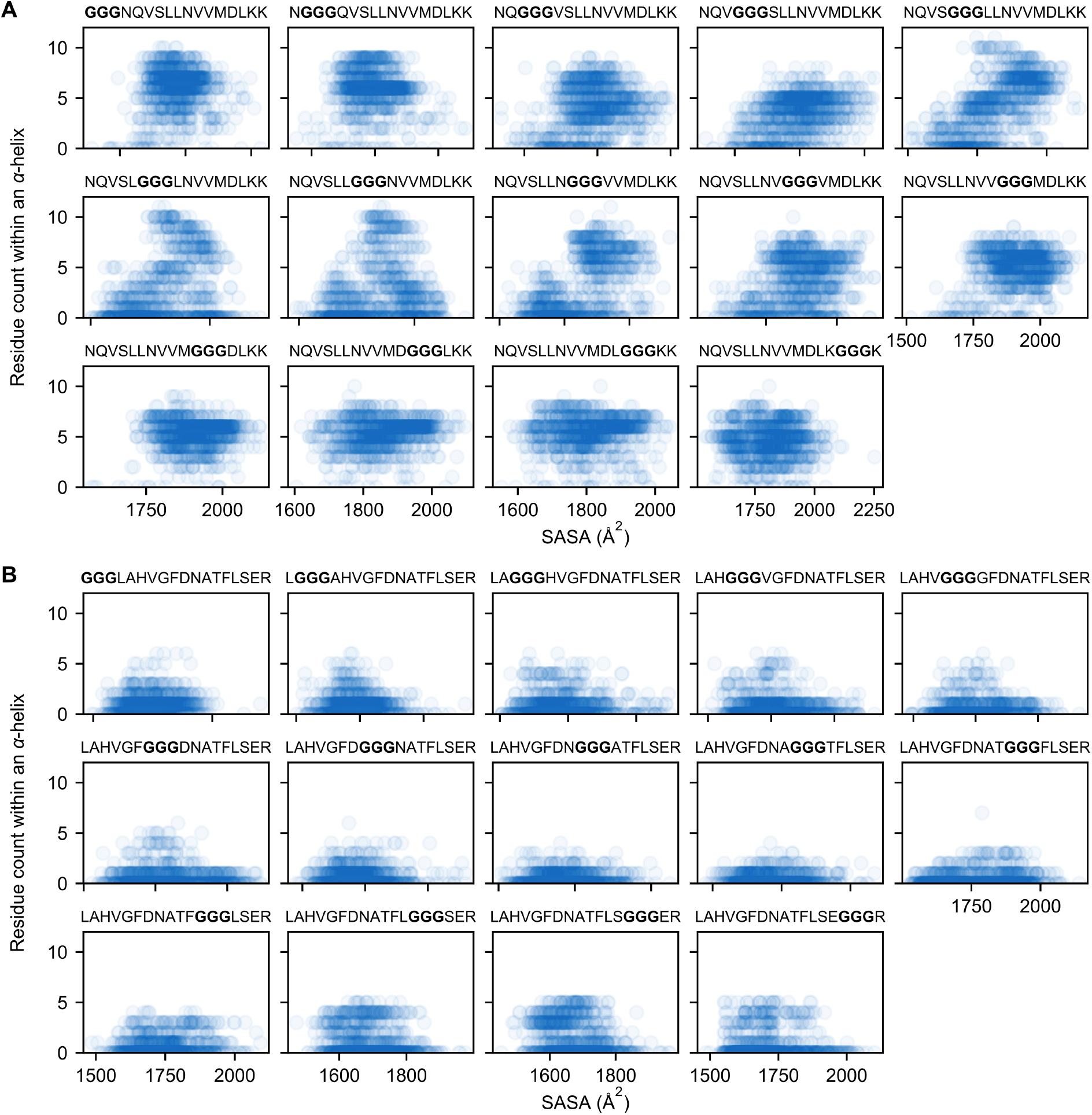
Residue count within an *α*-helix vs SASA. Scatter plot of the number of residues within an *α*-helix versus SASA across MD simulation runs, for each simulated peptide. Each plot contains 1000 points, one for each MD simulation run (*α* = 0.05). Plots corresponding to peptides in the high mode series grouped in (A), and those in the low mode series in (B). Glycine triplet is shown in bold.

